# Structural basis of Omicron neutralization by affinity-matured public antibodies

**DOI:** 10.1101/2022.01.03.474825

**Authors:** Daniel J. Sheward, Pradeepa Pushparaj, Hrishikesh Das, Changil Kim, Sungyong Kim, Leo Hanke, Robert Dyrdak, Gerald McInerney, Jan Albert, Ben Murrell, Gunilla B. Karlsson Hedestam, B. Martin Hällberg

## Abstract

The SARS-CoV-2 Omicron^1^ Variant of Concern (B.1.1.529) has spread rapidly in many countries. With a spike that is highly diverged from that of the pandemic founder, it escapes most available monoclonal antibody therapeutics^2,3^ and erodes vaccine protection^4^. A public class of IGHV3-53-using SARS-CoV-2 neutralizing antibodies^5,6^ typically fails to neutralize variants carrying mutations in the receptor-binding motif^7–11^, including Omicron. As antibodies from this class are likely elicited in most people following SARS-CoV-2 infection or vaccination, their subsequent affinity maturation is of particular interest. Here, we isolated IGHV3-53-using antibodies from an individual seven months after infection and identified several antibodies capable of broad and potent SARS-CoV-2 neutralization, extending to Omicron without loss of potency. By introducing select somatic hypermutations into a germline-reverted form of one such antibody, CAB-A17, we demonstrate the potential for commonly elicited antibodies to develop broad cross-neutralization through affinity maturation. Further, we resolved the structure of CAB-A17 Fab in complex with Omicron spike at an overall resolution of 2.6 Å by cryo-electron microscopy and defined the structural basis for this breadth. Thus, public SARS-CoV-2 neutralizing antibodies can, without modified spike vaccines, mature to cross-neutralize exceptionally antigenically diverged SARS-CoV-2 variants.

## INTRODUCTION

The Omicron variant harbors two deletions, one insertion, and 30 amino acid substitutions in the viral spike relative to the pandemic founder, including many in the receptor-binding domain (RBD). As a result of this antigenic drift, Omicron escapes the majority of monoclonal antibodies (mAbs), including those currently used in the clinic^2,3,12^ and displays markedly reduced sensitivity to neutralization by serum from convalescent and vaccinated individuals^3,13^. However, a third dose with vaccines based on the founder spike improved Omicron neutralization^14^, suggesting either the expansion of more cross-reactive B cells in the repertoire, the broadening of responses with affinity maturation, or both.

Several studies have characterized the B cell responses to SARS-CoV-2 in the context of infection and vaccination and revealed that the immunoglobulin heavy chain variable 3-53 gene (IGHV3-53) is the most frequently used by neutralizing SARS-CoV-2 antibodies^5,15^. These antibodies display convergent features and have previously been grouped into two ‘classes’ based on their targeted epitopes and characteristics^16^. “Class 1” antibodies are highly prevalent and use IGHV3-53 to bind to an epitope that overlaps the Receptor Binding Motif (RBM) that is only accessible in the RBD-up conformation. They typically possess short heavy chain complementarity-determining regions (HCDR3s) that make only minor contributions to the antibody’s interactions, and it is germline-encoded residues in IGHV3-53 that dominate their interaction, enabling potent neutralization of the founder spike with minimal affinity maturation^5^. However, antibodies of this class frequently fail to cross-neutralize variants carrying mutations in the RBM^7,8^.

Here, we describe a panel of IGHV3-53-using mAbs isolated from a convalescent individual approximately 7 months after infection. We show that a subset of these are broadly cross-neutralizing, capable of neutralizing Omicron and all other variants of concern (VoCs) tested, with exceptional potency. Additionally, we demonstrate the role of somatic hypermutation (SHM) at key positions in the acquisition of breadth. Further, we resolve a structure of the trimeric Omicron spike in complex with one of these broad mAbs, CAB-A17, at an overall 2.6 Å resolution, thereby defining the structural basis for cross-neutralization. These data demonstrate the potential for a common class of antibodies to develop broad cross-neutralization of SARS-CoV-2 through affinity maturation.

## RESULTS

### Identification of broadly cross-neutralizing IGHV3-53 antibodies

We expressed SARS-CoV-S spike-specific IGHV3-53 using mAbs (Fig. 1b) isolated from memory B cells from a convalescent individual, approximately seven months following an infection, and prior to vaccination.

**Figure 1.**
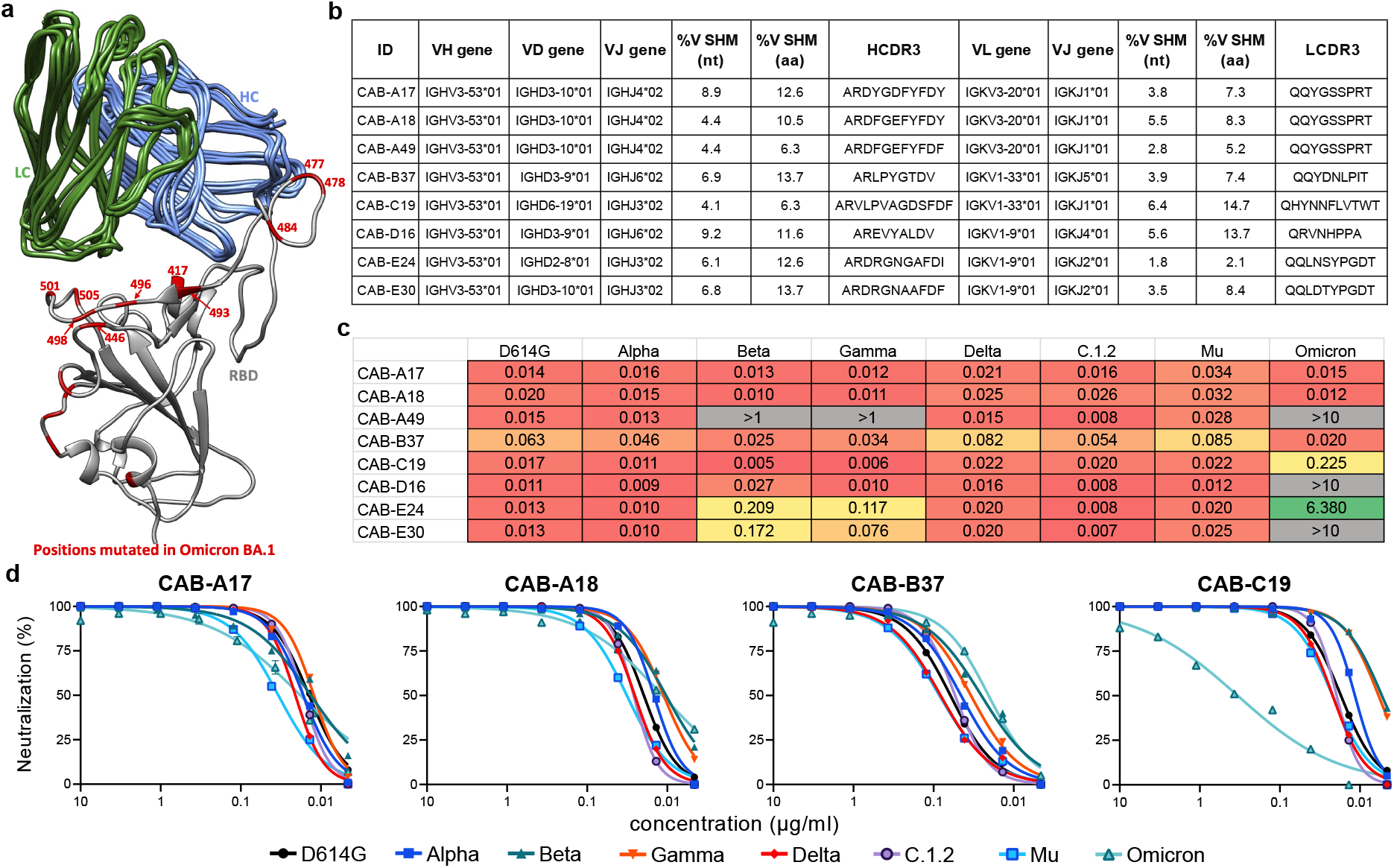
Identification of broadly cross-neutralizing SARS-CoV-2 antibodies. **a**. B.1.1.529 (Omicron) harbors multiple mutations at the interface with a public class of IGHV3-53-using neutralizing antibodies. The antibody heavy (HC) and light (LC) chains from 5 antibodies (P5A-3A1^18^, ab1^19^, COVOX-222^7^, BD-629^20^, C102^16^) are overlaid in complex with the SARS-CoV-2 RBD of the pandemic founder (grey) with the location of mutations in the Omicron RBD shown in red. **b**. Genetic features of the SARS-CoV-2 mAbs characterized here. **c**. Cross-neutralization of variants D614G (B.1), Alpha (B.1.1.7), Beta (B.1.351), Gamma (P.1), Delta (B.1.617.2), C.1.2, Mu (B.1.621), and Omicron (B.1.1.529) by isolated mAbs. Shown are the neutralizing IC_50_ titers (µg/ml). **d**. Neutralization curves against variants by the four mAbs able to cross-neutralize Omicron.

Using a pseudovirus neutralization assay, we investigated whether these mAbs could cross-neutralize both D614G and Omicron. While some showed significant reduction in potency against Omicron, three monoclonal antibodies from two clonal lineages exhibited no appreciable loss of potency against Omicron, which they neutralized with half-maximal inhibitory concentrations (IC50) <20 ng/ml (Fig. 1c and Fig. S1).

In addition to neutralizing Omicron, clonal lineages CAB-A (including mAbs CAB-A17 and CAB-A18) and CAB-B (with CAB-B37) exhibited exceptional breadth, potently neutralizing all other SARS-CoV-2 VoCs tested - including B.1.617.2 (Delta), B.1.351 (Beta), B.1.621 (Mu) and C.1.2 (Fig. 1d).

From a distinct lineage, CAB-C19 retained potency against all other VoCs, but lost around 20-fold potency against Omicron. Also from another lineage, CAB-D16 cross neutralized all other VoCs, but exhibited no neutralization of Omicron. From another lineage, CAB-E24 and CAB-E30 cross-neutralized Alpha, Delta, C.1.2 and Mu, but lost potency against Beta and Gamma, and showed little to no ability to neutralize Omicron. Like Omicron, the RBDs of Beta and Gamma are mutated at 417, which is crucial for antibodies of this class^9,10,17^. A similar neutralization pattern is shown by CAB-A49, which is from the otherwise-broad CAB-A lineage but with fewer amino-acid SHMs and retains potency against all VoCs except Beta, Gamma and Omicron, suggesting that specific maturational pathways are required for breadth.

### Affinity maturation towards cross-neutralization

While a germline-reverted version of CAB-A17 (glCAB-A17; with the mature CAB-A17 HCDR3) was capable of neutralizing D614G, albeit 55-fold less potently than the mature antibody, it had no detectable neutralization of Omicron (Fig. 2). To characterize the affinity maturation required for the development of cross-neutralization of Omicron by CAB-A17, we introduced select SHMs identified in the CAB-A17 lineage individually, or in combination, into glCAB-A17 and assessed their neutralization potency against D614G and Omicron using a pseudovirus assay. Mutations introduced individually enhanced the neutralization of D614G, but none afforded detectable cross-neutralization of Omicron (Fig 2b). However, the introduction of four mutations in the HC together: G26E, T28I, S53P, and Y58F afforded cross-neutralization of Omicron equivalent to the germline antibody against D614G and enabled potency against D614G approaching that of the mature antibody (Fig. 2b-d). S53P alone was sufficient to recapitulate these titers against D614G, but not against Omicron. However, T28I and S53P in combination on the background of the germline-reverted antibody enabled detectable cross-neutralization of Omicron (Fig. 2b-d). Together, these data highlight that relatively few somatic hypermutations can confer breadth to a commonly-elicited class of IGHV3-53 antibodies, even extending to the highly divergent Omicron RBM.

**Figure 2.**
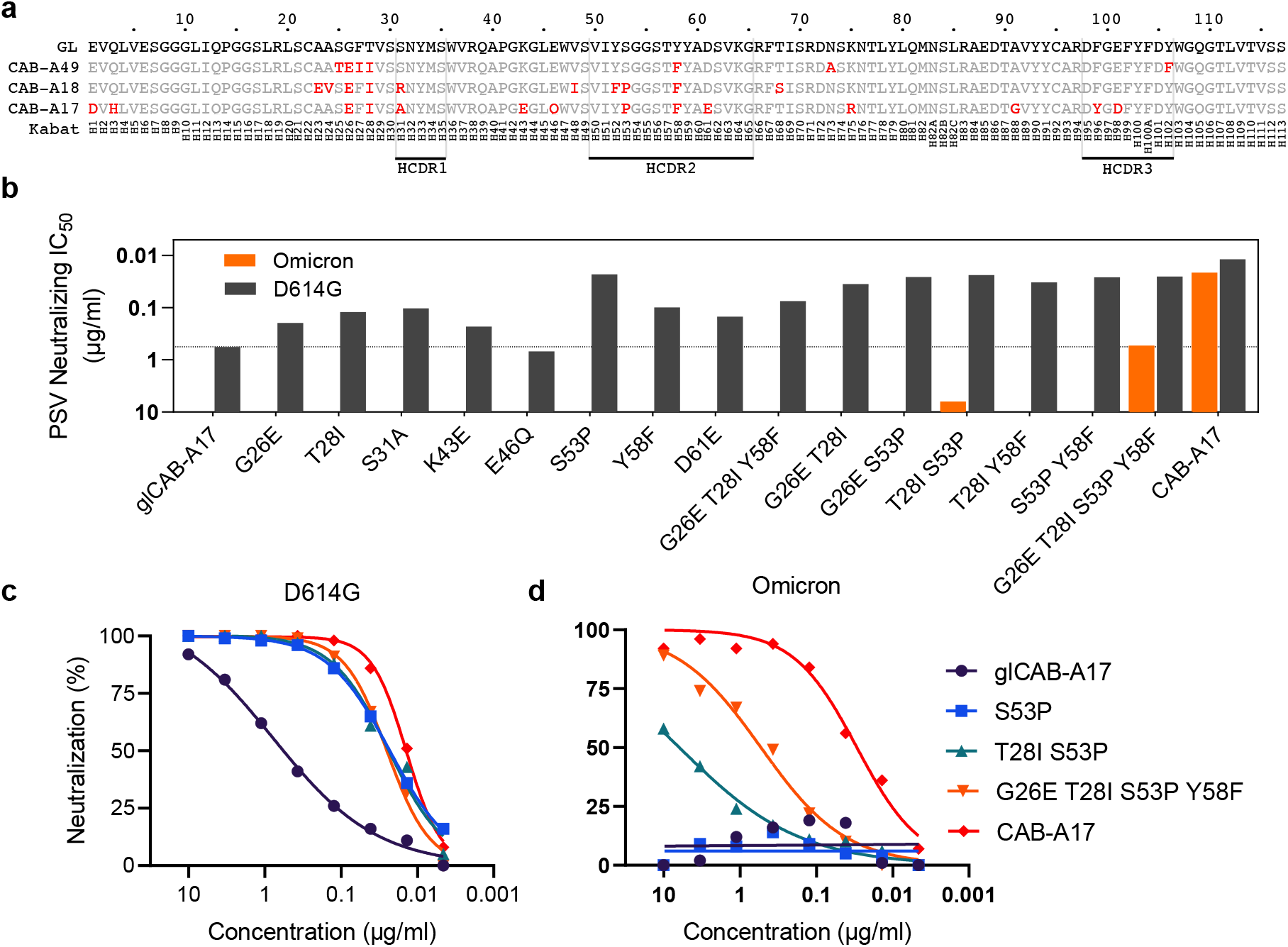
Affinity maturation enables cross-neutralization of Omicron. **a**. Alignment of heavy chain sequences of mAbs from lineage CAB-A. CDR borders and numbering are shown according to the Kabat scheme. Somatic hypermutations away from germline (GL) are shown in red. **b**. The effect of the introduction of identified SHMs, individually or in combination, into the glCAB-A17 Heavy Chain on the neutralization of D614G (black) and Omicron (Orange). **c-d**. Neutralization of **c**. D614G and **d**. Omicron by CAB-A17, and the germline-reverted version of CAB-A17 (glCAB-A17) as well as versions carrying introduced SHMs.

### Cryo-EM structure of CAB-A17 in complex with the Omicron spike reveals the structural basis for broad neutralization

To define the mode of recognition permitting such broad and potent neutralization, we resolved the structure of a CAB-A17 Fab in complex with the Omicron spike. The cryo-EM reconstruction, with an overall resolution of 2.6 Å, revealed a 2-up conformation of the spike and each up-RBD was decorated with a CAB-A17 Fab bound in the ACE2 binding surface (Fig. 3a-b). All spikes in the cryo-EM reconstruction were in the 2-up conformation and bound two Fabs, albeit with different conformations. To obtain a molecular understanding of the CAB-A17-Omicron RBD interaction, we utilized localized-reconstruction techniques on a subset of the particles. The observed CAB-A17 Fab binding mode is compatible with bivalent binding of full IgG to the Omicron spike. The interface area from the heavy-chain interaction (829 Å^2^. was towards the higher end of structurally determined representatives of IGHV3-53 derived SARS-CoV-2 RBD binders^17^. In contrast, the light chain interaction area (310 Å^2^. was more typical. To understand the molecular basis for the acquisition of breadth by this class of IGHV3-53-using antibodies, we analyzed the interaction between the RBD and the heavy and light chains of CAB-A17. All CDRs of the light and heavy chains of CAB-A17 interacted with Omicron RBD (Fig. 3c) through surface complementarity, residue packing interactions and specific hydrogen bonds (Table S1). Remarkably, seven of the interaction-interface residues on the Omicron spike are mutated relative to the founder variant (Fig. 3d). Like the other representatives of the public class of IGHV3-53-using neutralizing antibodies, CAB-A17 binds in or in the immediate vicinity to the ACE2-RBD interface (Fig. 3e and 4a).

**Figure 3.**
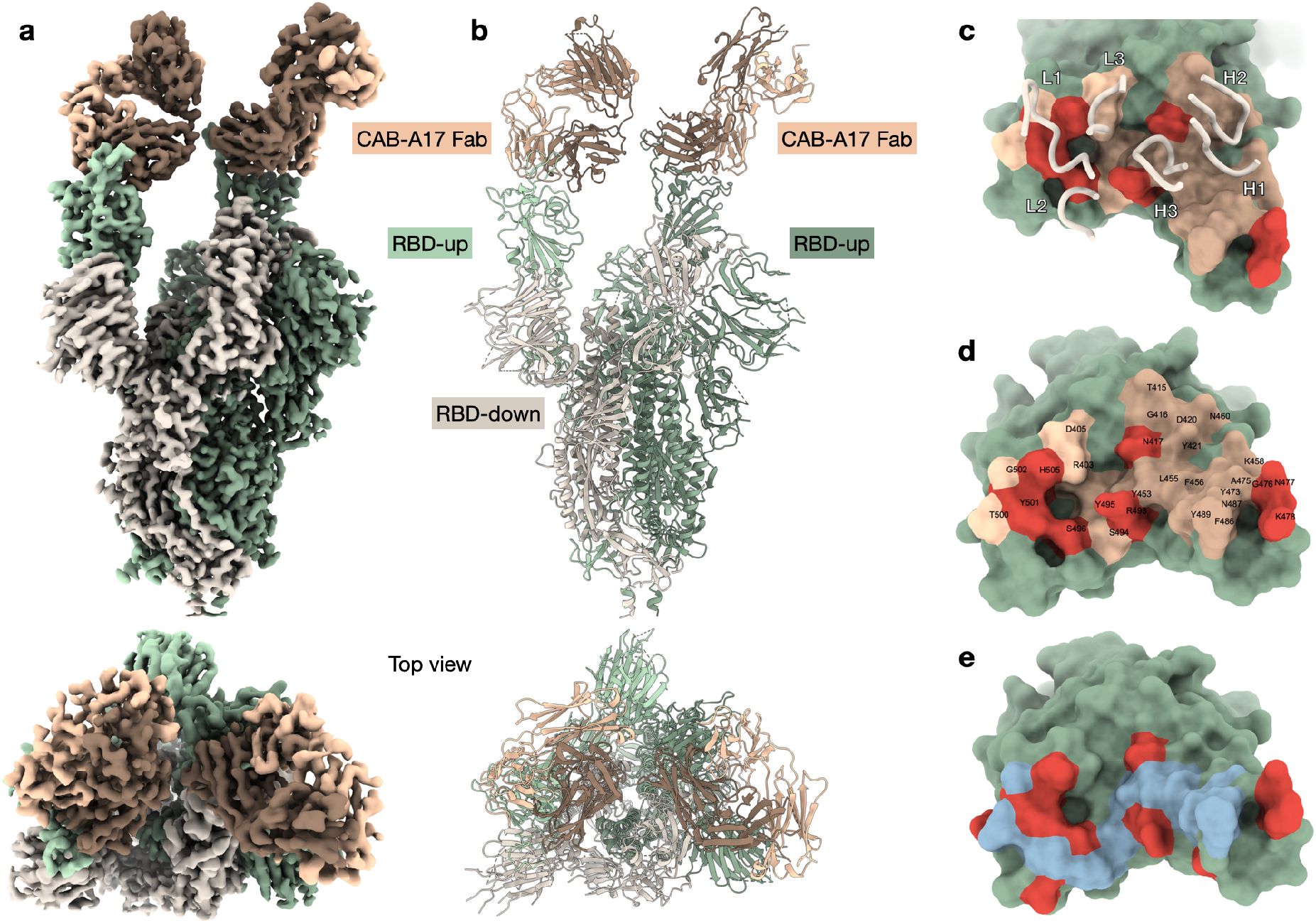
Cryo-EM reconstruction of the CAB-A17 Omicron-Spike complex. **a**. Cryo-EM reconstruction of CAB-A17 Fab bound to the Omicron spike determined to an overall resolution of 2.6 Å. For illustration and clarity purposes, the map is a composite of the local reconstruction (3.6 Å) and the overall reconstruction. Protomers in RBD-up conformation are colored green and light green. The RBD-down conformation protomer is colored in beige. The CAB-A17 Fab light and heavy chains are colored in light and dark brown, respectively. **b**. Cartoon representation of the molecular structure with the same coloring as in a. **c**. The epitope of CAB-A17 mapped on the Omicron RBD interface. Light and heavy CDR loops are labeled L1–L3 and H1–H3, respectively. Epitope residues contacting the light and heavy chains are shown in light brown and dark brown, respectively. Residues that are mutated in the Omicron variant are labeled red-orange. **d**. Epitope residues labeled. The same coloring as in **c. e**. ACE2-Omicron RBD interface with ACE2 contacting residues labeled in light blue and residues mutated in Omicron labeled red-orange.

**Figure 4.**
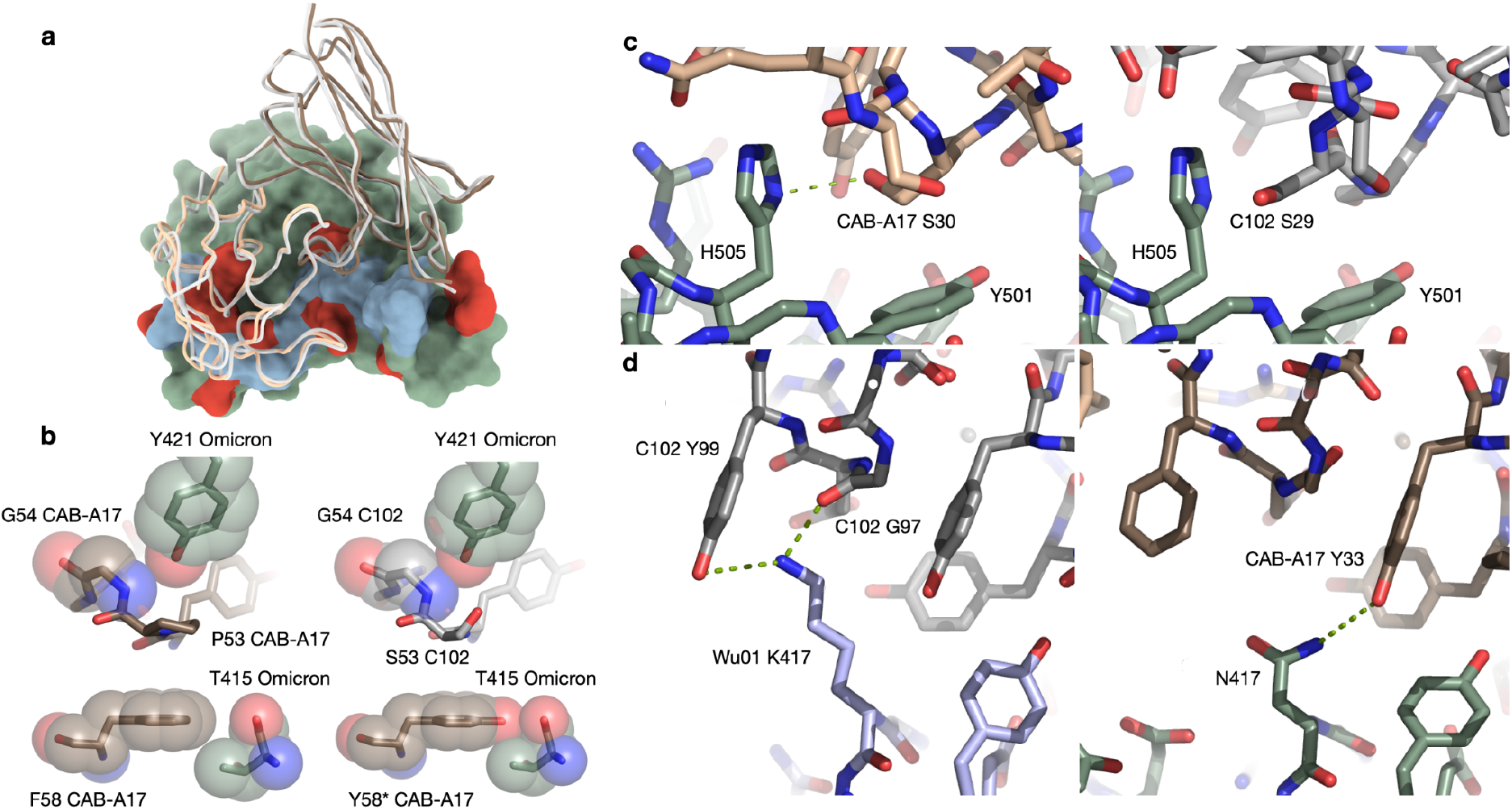
Detailed comparison between CAB-A17 and C102. **a**. Superimposition of CAB-A17 and C102. C102 is colored in light gray. The CAB-A17 Fab light and heavy chains are colored in light and dark brown, respectively. **b**. Modeling of the interaction between Y421 and P53/S53/G54 in the HCDR2. Relevant atoms are shown in both stick and in transparent VDW-sphere representation. **c**. Interactions in the LCDR1. CAB-A17 S30 (left) forms a hydrogen bond with H505, while S29 in C102 (right) clashes into H505. **d**. K417 in the founder variant (Wu01) forms two strong hydrogen bonds to the C102 G97 and Y99 (left), while these are lost in Omicron (right). In contrast, the N417 in Omicron can form a hydrogen bond with CAB-A17 Y33.

As the interaction between the SARS-CoV-2 founder RBD and IGHV3-53-using class 1 antibodies is well-explored^17^, we focused on interactions enabling cross-neutralization of Omicron by CAB-A17 by comparison with the mAb C102^16^. C102 and CAB-A17 are both class 1 antibodies^16^, and share an overall binding mode^21^ to an epitope overlapping the ACE2 interaction surface (Fig. 4a), with a C_α_-RMSD of 0.62 Å^2^ when aligned to their respective RBDs. C102 uses the same light chain gene (IGKV3-20), and has 91% amino acid identity to the CAB-A17 heavy chain (Fig. S4). However, while C102 potently neutralizes the SARS-CoV-2 founder and Delta (IC50 52 ng/ml and 17 ng/ml, respectively), it fails to neutralize Beta and Omicron (IC50 > 10 μg/ml)^11^, indicating that key parts of the C102 binding interface are affected by mutations in Omicron.

Pseudovirus neutralization assays showed that an S53P in the HCDR2 is beneficial for neutralization of D614G, and enables, but alone is insufficient for Omicron neutralization (Fig. 2b). In C102, the S53 remains in the germline configuration, thereby allowing a structural comparison of this change. In both C102 and CAB-A17, the heavy chain backbone changes direction 180 degrees, and this sharp turn enables binding of RBD Y421 in a pocket formed by the turn and to hydrogen bond to the backbone amide of G54, and in C102, also the S55 hydroxyl. However, in the Omicron RBD, an apparent backbone shift pushes the RBD Y421 hydroxyl such that it would clash with G54, while in CAB-A17 the slightly sharper turn facilitated by P53 retains the favorable interaction and conformation for RBD Y421 despite the backbone shift (Fig. 4b-c). Interestingly, the S53P change occurs preferentially with Y58F (Supp. Section “Tabulating SHM frequencies”). Modeling an F58Y mutation in the CAB-A17 structure shows that the hydroxyl of a 58Y would clash into RBD T415 (Fig. 4B). Hence, S53P and a Y58F may act in concert to retain a favorable interaction with RBD Y421. In addition to the S53P and Y58F pair, combined reversion together with G26E and T28I enabled a partial recovery of neutralization capacity against Omicron (Fig. 2d). E26 in CAB-A17 has the potential to contact K478 (T478 in the founder variant), thereby promoting RBD interaction. Nevertheless, even without a T478K mutation, any change from a G26 would stabilize the HC interaction with the RBD region 475-478.

In the CAB-A17 light-chain, SHMs in the LCDR1, including a 1 AA deletion, changed the VSSS-motif to an LST (Fig. S4). This led to a slight shortening of the backbone in a critical region close to the mutated Q498R and N501Y in the LC interaction area (Fig. 4c). This shortening enabled CAB-A17 to retain a favorable RBD binding mode (including a hydrogen bond with H505) (Fig. 4c and Table S1) despite many mutations in this part of the interface. However, even without the LCDR1 deletion, binding in this region may still occur, but with a change in Ab-RBD approach angle. This may explain the lack of full restoration of neutralization activity even after the re-introduction of four key heavy chain SHMs into glCAB-A17.

A key mutation for viral escape from IGHV3-53 public antibodies is K417N, first described in the Beta variant^17^. The replacement of K with N in Omicron removes two strong hydrogen bonds from the interaction between the C102 LCDR3 (Y99 and G97) and the founder variant RBD. In contrast, a new hydrogen bond, albeit long, is formed between Omicron N417 and Y33 in the CAB-A17 LCDR1 (Fig. 4d).

## DISCUSSION

On aggregate, Omicron exhibits the most extensive resistance to neutralizing antibodies of all SARS-CoV-2 variants that have emerged so far^2,3,12,13^. To date, only a few monoclonal antibodies have been confirmed to cross-neutralize Omicron. This includes S309/sotrovimab^22^ targeting a conserved epitope on the side of the RBD, but also three antibodies targeting the RBM^23^. Clinically-approved/authorized monoclonal antibodies casirivimab and imdevimab (Regeneron), bamlanivimab and etesevimab (Eli Lilly) show no detectable neutralization of Omicron^2,3,24^, with tixagevimab and cilgavimab (AstraZeneca) showing low but detectable anti-Omicron neutralization^2,24^.

Here, we describe several extremely potent neutralizing antibodies, capable of neutralizing all SARS-CoV-2 variants of concern evaluated, including Omicron. In our hands, CAB-A17 is 15-fold more potent against Omicron than S309 ^3^, which is currently the only clinically-approved mAb with retained activity. CAB-A17 (and clonally-related CAB-A18) use IGHV3-53 and IGHJ4, have a short (11 AA) HCDR3 and are paired with IGKV3-20, all typical of common “class 1”^16^ antibodies (public clonotype 2^15^).

The IGHV3-53 gene is frequently used in the human antibody repertoire. All currently known alleles of this gene contain germline-encoded motifs in the HCDR1 and HCDR2 (NY and SGGS, respectively) that enable their interaction with the class 1 epitope in the SARS-CoV-2 RBM. The closely related IGHV3-66 gene similarly uses these motifs^15,17^. IGHV3-53 is highly overrepresented amongst potent SARS-CoV-2 neutralizing monoclonal antibodies^5,25–27^. The precursor frequency of this class of neutralizing antibodies in the human repertoire was estimated to be around 1: 44,000^28^, suggesting that such antibodies are readily elicited in most people.

CAB-A17 and CAB-A18 here are more affinity matured (V gene SHM: 12.6% AA, 8.9% nuc, and 10.5% AA, 4.4% nuc, respectively) than the previously described prototypical members of this class - likely reflecting the relatively delayed isolation, seven months following a confirmed SARS-CoV-2 infection. CAB-A49 from the same lineage but with fewer amino-acid SHMs (6.3%) did not exhibit the same breadth, suggesting the importance of somatic hypermutation in the development of breadth. However, only four SHMs in the HC alone were required to neutralize Omicron with the same potency as the germline antibody against D614G, indicating the mutational barrier to breadth is not prohibitive. This suggests that specific but not extensive SHM pathways are required for breadth by this antibody class.

Importantly, the mAbs described here were isolated in Sweden in late 2020, before VoCs circulated in the region^29^. This is encouraging, as it indicates that mutations selected for in the affinity maturation to the founder spike also afforded improved cross-neutralization of Omicron. The detection of cross-neutralizing antibodies within the repertoire of a convalescent individual also suggests that such memory B cells are available to be recruited upon re-infection or vaccination.

Here, a cryo-EM structure of CAB-A17 in complex with Omicron spike reveals the structural basis of this broad cross-neutralization. This is enabled, in part, by SHMs including S53P and Y58F, which allow favorable interactions despite a structural shift in Omicron that would otherwise potentially lead to a clash between S53 in the antibody HCDR2 and the tyrosine at spike position 421.

In all germline V3-53 and V3-66 alleles, the serine at position 53 is encoded by the codon AGC, which requires at least two nucleotide changes to mutate to a proline. This mutational barrier likely explains why the S53P mutation, which offers the largest improvement in neutralization potency against D614G of all single mutations we tested, is not more frequently observed.

IGHV3-53 using mAbs isolated to date frequently fail to neutralize Beta, and Gamma^7^, or K417N/T mutants^8,10,17^. Three of our antibodies in this class appear to be affected by K417N/T and lose potency to Beta, Gamma and Omicron. This includes CAB-A49 from the same clonal lineage as the ultra-broad CAB-A17 and CAB-A18. Two other mAbs (CAB-C19 and CAB-D16) that are resilient to mutations at 417, neutralizing Beta and Gamma, still lose potency, to varying degrees, against Omicron. These are reminiscent of one previously published antibody from this class, COVOX-222, which exhibits significant breadth against Alpha, Beta, Gamma and Delta^7^, but loses potency against Omicron by around 13-fold^30^.

Light chain interactions also play a critical role in the neutralization of variants by this class of antibody, where the LCDR1 of these antibodies make direct contacts with the spike near residue 501, though these can vary substantially^5^. In line with this, the light chain from COVOX-222 could rescue neutralization of Beta and Gamma by other IGHV3-53 antibodies^7^. Omicron carries not only N501Y but also proximal mutations at 496, 498, and 505. Here, we show that CAB-A17 has a single amino acid deletion, shortening the LCDR1 in a critical region close to this interface. The clonally-related CAB-A18 does not have this shortening but carries other mutations in the LCDR1, suggesting that multiple pathways to avoid clashes exist.

In a relatively small set of antibodies from this class, we found three antibodies from two clonally distinct lineages that could cross-neutralize all tested variants, including Omicron. The sequences are sufficiently different between these broad neutralizers to suggest that there are distinct ways of overcoming the mutations in Omicron and other variants, and achieving this degree of breadth and potency. The breadth and potency of these antibodies make them important therapeutic candidates in the context of an antigenically evolving pandemic. The identification of multiple broadly cross-neutralizing antibodies from this public class, capable of neutralizing Omicron without loss of potency, also suggests that these responses may be more generally achievable than previously appreciated.

There is increasing evidence that an additional dose of one of the licensed SARS-CoV-2 vaccines enables substantial improvements in the cross-neutralization of Omicron^14^, suggesting that affinity maturation may broaden responses. Indeed, affinity maturation of antibody lineages occurs over the course of months after SARS-CoV-2 infection, and is associated with the cross-neutralization of variants of concern^31^. Here, we demonstrate the potential for a common class of public antibodies to develop broad cross-neutralization through affinity maturation, without modified spike vaccines.

## Acknowledgements

We acknowledge the G2P-UK National Virology consortium funded by MRC/UKRI (grant ref: MR/W005611/1.) and the Barclay Lab at Imperial College for providing B.1, B.1.1.7, B.1.351, P.1, B.1.617.2, B.1.621 and C.1.2 spike plasmids. pCMV-dR8.2 dvpr was a gift from Bob Weinberg (Addgene plasmid # 8455; http://n2t.net/addgene:8455; RRID:Addgene_8455). pBOBI-FLuc (Addgene plasmid # 170674; http://n2t.net/addgene:170674; RRID:Addgene_170674), and pcDNA3.3_CoV1_D28 (Addgene plasmid # 170447 ; http://n2t.net/addgene:170447 ; RRID:Addgene_170447) were gifts from David Nemazee. We further acknowledge the Karolinska Institutet’s 3D-EM facility, which was used for collection of all cryo-EM data.

## Author contributions

Conceptualization: DJS, PP, BM, GBKH, BMH

Methodology: DJS, PP, LH, BM, GBKH, BHM

Investigation: DJS, PP, HD, CK, SK, BMH

Visualization: DJS, PP, HD, BM, BMH

Resources: LH, RD, GM, JA

Funding acquisition: GM, GBKH, BM, BMH

Supervision: BM, GBKH, BMH

Writing – original draft: DJS, BM, GBKH, BMH

Writing – review & editing: DJS, PP, HD, BM, GBKH, BMH

## Data and materials availability

The cryo-EM reconstructions have been deposited in the Electron Microscopy Data Bank under accession codes EMD-xxxxx (Omicron spike + Fab fragment of CAB-A17), EMD-xxxxx (localized reconstruction of Omicron RBD + Fab fragment of CAB-A17). The atomic coordinates have been deposited in the Protein Data Bank under IDs xxxx (Omicron spike + Fab fragment of CAB-A17), xxxx (localized reconstruction of Omicron RBD + Fab fragment of CAB-A17). Nucleotide sequences for antibodies described in this publication are available at GenBank, with accessions XXXXXXXX-XXXXXXXX. All other datasets generated during and/or analyzed during the current study are available from the corresponding authors on reasonable request.

## METHODS

### Sample collection

Hospital workers at the Karolinska University Hospital in Stockholm, Sweden, were invited to participate in a study that aimed to characterize their antibody responses following SARS-CoV-2 infection. Participants who were confirmed PCR-positive in May 2020 provided blood samples in December 2020. Informed consent was obtained from all participants as part of ethics approvals (Decisions# 2020-01620, 2020-02881 and 2020-05630) from the National Ethical Review Agency of Sweden. Blood was collected in EDTA tubes for the separation of plasma and peripheral blood mononuclear cells (PBMCs). PBMC isolation was performed by density-gradient centrifugation using Ficoll-Paque PLUS (GE Healthcare) or SepMate™ PBMC Isolation tubes (Stem Cell technologies). After a gradient separation the mononuclear cell layers were collected and the cell pellets were washed with sterile PBS. The isolated PBMCs were frozen in FBS with 10% dimethyl sulfoxide (DMSO) (Sigma).

### Single-cell sorting of SARS-CoV-2-specific memory B cells by flow cytometry

Spike-specific memory B cells were stained and single-cell-sorted from PBMCs using a three-laser FACSAria cell sorter (BD Bioscience) by gating on live, CD3-, CD14-, CD20+, CD27+, IgG+, spike+ cells into 96-well PCR plates containing 4 μl of cell lysis buffer. All fluorescently labeled antibodies and the biotinylated spike-trimer conjugated to streptavidin-allophycocyanin (SA-APC) (Invitrogen) were titrated before sorting. Plates were sealed and immediately frozen on dry ice and stored at -80 °C.

### Single B cell RT-PCR

The 96-well plates containing lysed single B cells were thawed on ice and cDNA was generated by reverse transcription using random hexamers, dNTPs and SuperScript IV reverse transcriptase (Invitrogen). V(D)J sequences were amplified separately in 25 µl nested PCR reactions using 3 µl of cDNA in the 1^st^ round PCR and 1.5 µl PCR product in the 2^nd^ round PCR. The HotStarTaq Plus Kit (Qiagen) and 5’ leader sequence-specific and 3’ IgG-specific primers were used. PCR products from positive wells were purified, Sanger sequenced (Genewiz) and analyzed.

### Cloning of monoclonal Abs

HC and LC sequences were cloned into expression vectors containing human Igγ1 H, Igκ1 L, or Igλ2 L constant regions^32^ by Gibson assembly. Briefly, HC and LC V(D)J sequences were synthesized with overhangs that match the ends of the linearized vectors. Gibson assembly was performed using a Gibson master mix (New England Biolabs), 50ng of vector, 30 ng of insert in a 20ul reaction, incubated at 50°C for 1 hour, and transformed into XL10-Gold ultracompetent cells by heat shock at 42 °C for 30 seconds, according to the manufacturer’s protocol (Agilent Technologies). Colonies were screened by PCR and confirmed by Sanger sequencing (Genewiz). Positive colonies were expanded and plasmids purified using a plasmid Midi-prep kit (Qiagen) as per the manufacturer’s instructions.

### Expression and purification

For mAb expression, 15 μg of each HC and LC vector DNA was transfected into FreeStyle 293-F cells, cultured in 30 ml of FreeStyle 293 expression medium (Life Technologies) at a cell density of 1 × 10^6^ cells/ml and ≥ 90% viability, using 30 μl of 293fectin (Life Technologies) according to the manufacturer’s protocol. mAbs were harvested and purified 7 days after transfection using Protein G Sepharose columns (GE Healthcare). All purified recombinant mAbs were analyzed by SDS-PAGE under reducing conditions using NuPAGE Novex 4-12% Bis-Tris polyacrylamide gels and NuPAGE reducing agent (Life Technologies) according to the manufacturer’s instructions and by binding to SARS-CoV-2 spike and RBD by ELISA.

### Cell culture

HEK293T cells (ATCC CRL-3216) and HEK293T-ACE2 cells (stably expressing human ACE2) were cultured in Dulbecco’s Modified Eagle Medium (high glucose, with sodium pyruvate) supplemented with 10% fetal calf serum, 100 units/ml penicillin, and 100 μg/ml streptomycin. Cultures were maintained in a humidified 37°C incubator (5% CO_2_).

### Spike expression vectors

Spike expression vectors for pseudovirus neutralization assays encoded the variant spikes with the last 19 amino acids (or 28 amino acids in the case of SARS-CoV) truncated to enhance incorporation. SARS-CoV-2 B.1, B.1.1.7, B.1.351, P.1, B.1.617.2, B.1.621 and C.1.2 spike expression plasmids were obtained from the G2P-UK National Virology consortium and the Barclay Lab (Imperial College). The B.1.1.529 spike was molecularly cloned from an infected individual^3^. The SARS-CoV spike expression vector was a gift from David Nemazee (Addgene #170447).

### Pseudovirus Neutralization Assay

Spike-pseudotyped lentivirus particles were generated by the co-transfection of HEK293T cells with a spike-encoding plasmid, an HIV gag-pol packaging plasmid (Addgene #8455), and a lentiviral transfer plasmid encoding firefly luciferase (Addgene #170674) using polyethylenimine (PEI). Pseudovirus neutralization assays were performed using HEK293T-ACE2 cells, as previously described^29^. Briefly, pseudoviruses were incubated with serial 3-fold dilutions of serum for 60 minutes at 37°C in black-walled 96-well plates. 10,000 HEK293T-ACE2 cells were then added to each well, and plates were incubated for 44-48 hours. Luminescence was measured using Bright-Glo (Promega) on a GloMax Navigator Luminometer (Promega). Neutralization was calculated relative to the average of 8 control wells infected in the absence of serum. All fold-change comparisons used IC50 values from neutralization assays run side-by-side.

### Tabulating SHM frequencies

From the Coronavirus-Binding Antibody Sequences & Structures (CoV-AbDab) database^33^, we identified 162 anti-SARS-CoV-2 RBD monoclonal antibodies that were of human origin, used V3-53 or V3-66 germlines, had a CDR3 length < 13, and used either V3-20, V1-9, or V1-33 light chain V genes. Of these, 9% have G26E, 25% have T28I, 10% have S53P, and 58% have Y58F. 53P occurs 13 times with 58F and only 4 times with Y58.

### Preparation of Fab fragments

Fab fragments were prepared by digesting IgG with immobilized papain and separation of Fab and Fc fragments with a Protein A column, using the Pierce Fab Preparation Kit (ThermoFisher Scientific) per the manufacturer’s instructions.

### Cryo-EM sample preparation and imaging

Spike trimer (1.5 mg/ml; B.1.1.529 Omicron spike HexaPro^34^) and a Fab preparation of CAB-A17 were mixed in a 1:0.8 molar ratio (S protein monomer–Fab), followed by incubation on ice for 20 min. Prior to cryo-EM grid preparation, grids were glow-discharged with 25 mA for 2 min using an EMS 100X (Electron Microscopy Sciences) glow-discharge unit. Grids used were CryoMatrix holey grids with amorphous alloy film (Nitinol; R 2/1 geometry; Zhenjiang Lehua Technology Co., Ltd). 3-μl aliquots of Fab preparations were applied to the grids, grids were then hand-blotted from the side followed by two applications of the pre-incubated Omicron-Fab mixtures with intermittent hand-blotting. Finally, the grids with sample were vitrified in a Vitrobot Mk IV (Thermo Fisher Scientific) at 4°C and 100% humidity [blot 11 s, blot force 4, 595 filter paper (Ted Pella Inc.)].

Cryo-EM data collection was performed using EPU 2.13 (Thermo Fisher Scientific) using a Krios G3i transmission-electron microscope (Thermo Fisher Scientific) operated at 300 kV in the Karolinska Institutet’s 3D-EM facility. Movies were acquired in nanoprobe EFTEM SA mode at 165 kx nominal magnification with a slit width of 10 eV using a K3 Bioquantum (operated in CDS mode) for 2 s during which 60 movie frames were collected with a fluency of 0.91 e^™^/Å^2^ per frame (see table S2). A stage tilt of 20-degrees was used to alleviate preferred particle orientations. Motion correction, and Fourier cropping (to 1.01 Å per pixel), were performed on the fly using Warp^35^. Micrographs were imported into CryoSPARC 3.31 ^36^ and CTF parameters were estimated using a patch model.

A total of 15,106 micrographs were selected on the basis of an estimated resolution cutoff of 6 Å and defocus below 4 microns. Particles were picked by CryoSPARC through template matching using projections generated from a previous reconstruction of the ancestral SARS-CoV-2 spike. Extracted particles were used for a four-class ab-initio model generation followed by three rounds of heterogeneous refinement in cryoSPARC with ensuing nonuniform 3D refinement of the resulting particles for the spike containing particle set. Local CTF refinements were performed interspersed with global aberration estimation and correction (beam tilt, trefoil, tetrafoil and anisotropic magnification). The two bound Fabs of CAB-A17 had varying orientations relative to the body of the spike and relative to each other. Therefore, 3D-classification was performed within CryoSparc 3.31 without pose alignment using 20 classes with a mask around the two RBDs in up conformations and their bound Fabs. One class was selected for further processing and from the particles (57,905 out of a total of 398,386) belonging to this class, we performed local reconstruction of the volume close to one of the RBD-Fab interfaces. This process significantly enhanced the resolvability of the map and thereby enabled molecular fitting and interpretation of the maps. All particles were processed with C1 symmetry. Please see table S2 for data collection, processing statistics.

### Cryo-EM model building and structure refinement

The structure of the ancestral spike protein trimer in 2-up conformation PDB: 7A29^37^ was used as a starting model for model building. The model was mutated and rebuilt manually to reflect the Omicron spike sequence and structure. Structure refinement and manual model building were performed using COOT^38^ and PHENIX^39^, respectively, in interspersed cycles with secondary structure, Ramachandran, rotamers and bond geometry restraints. Structure figures and EM density-map figures were generated with UCSF ChimeraX^40^ and COOT, respectively. Please see table S2 for refinement and validation results.

### Statistical analysis

Neutralizing IC50 values were calculated in Prism v9 (GraphPad Software) by fitting a four-parameter logistic curve, to neutralization by serial 3-fold dilutions. Neutralization was bounded between 0 and 100%.

## Supplementary Information

**Fig. S1.**
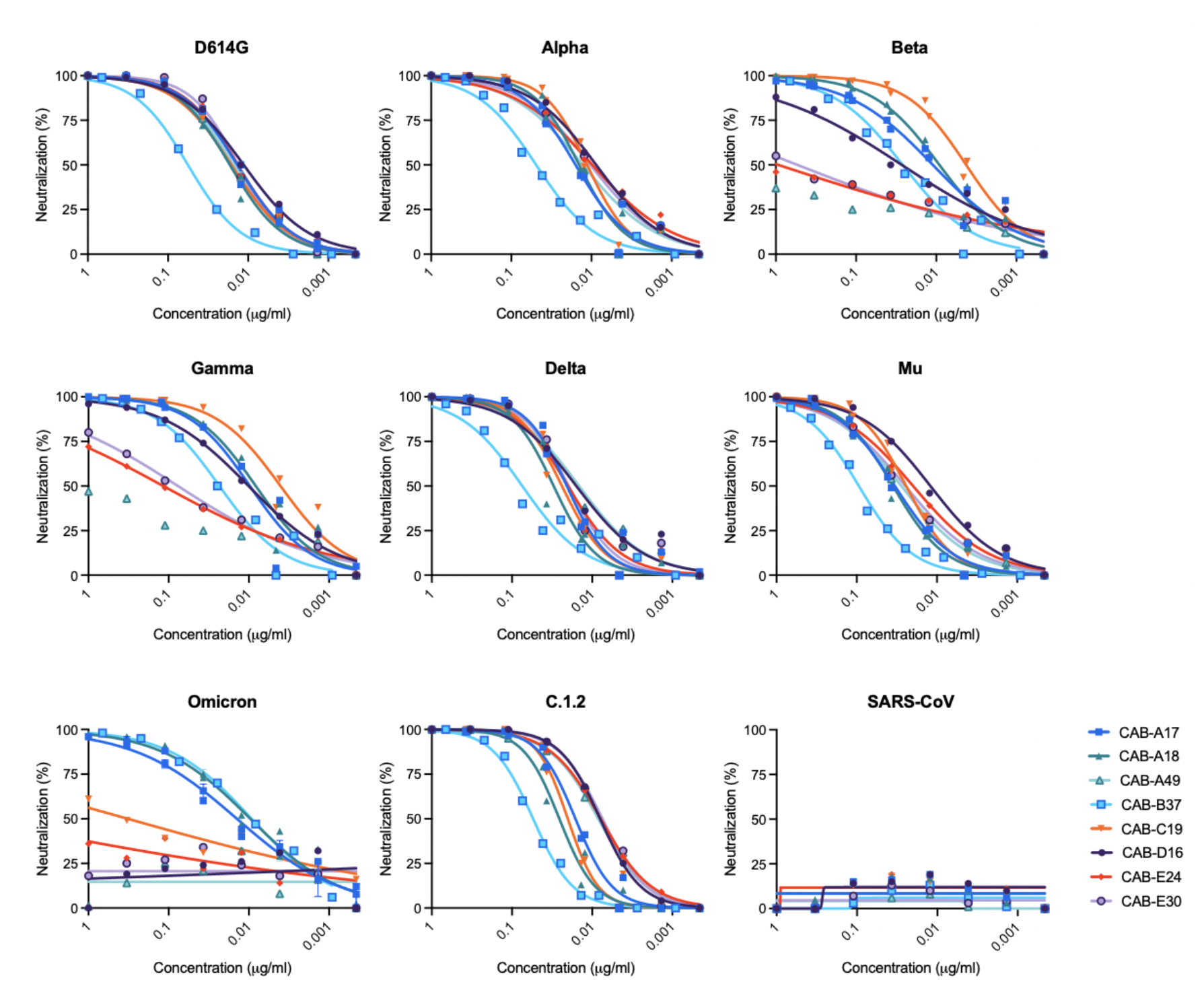
Broad and potent neutralization by IGHV3-53 antibodies isolated from the memory B cell repertoire of a convalescent individual.

**Fig. S2.**
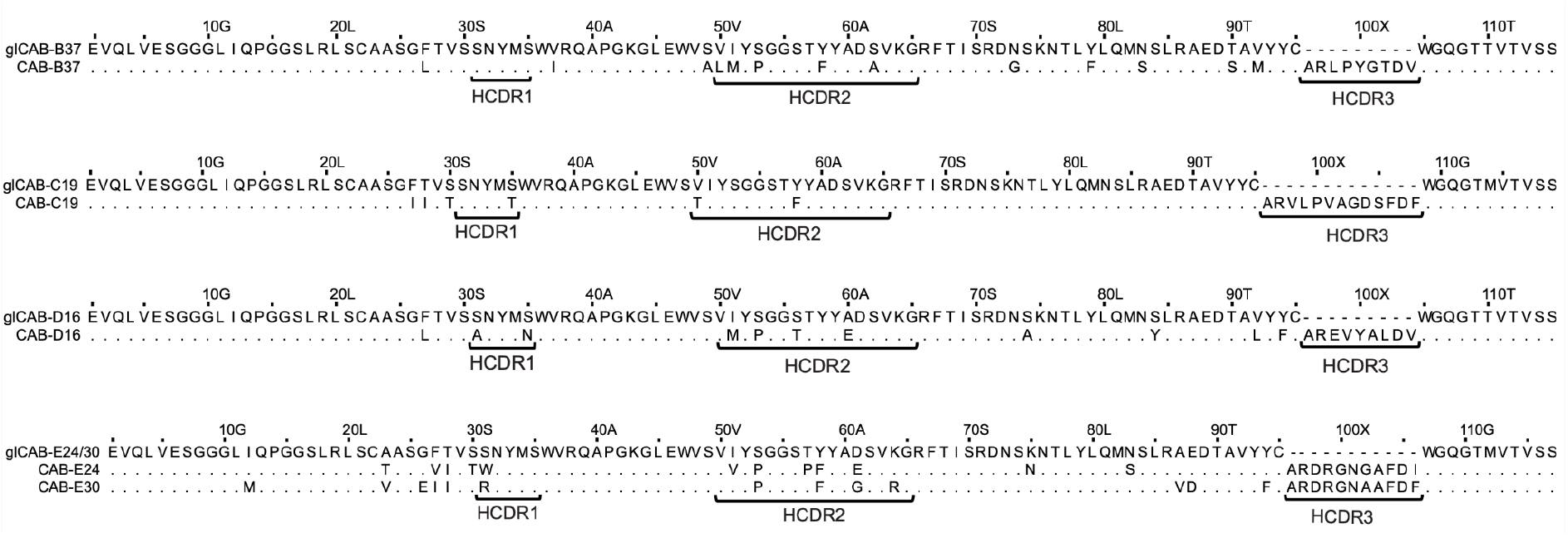
Alignment of heavy chain sequences with their inferred germline for four additional antibody lineages characterized here.

**Fig. S3.**
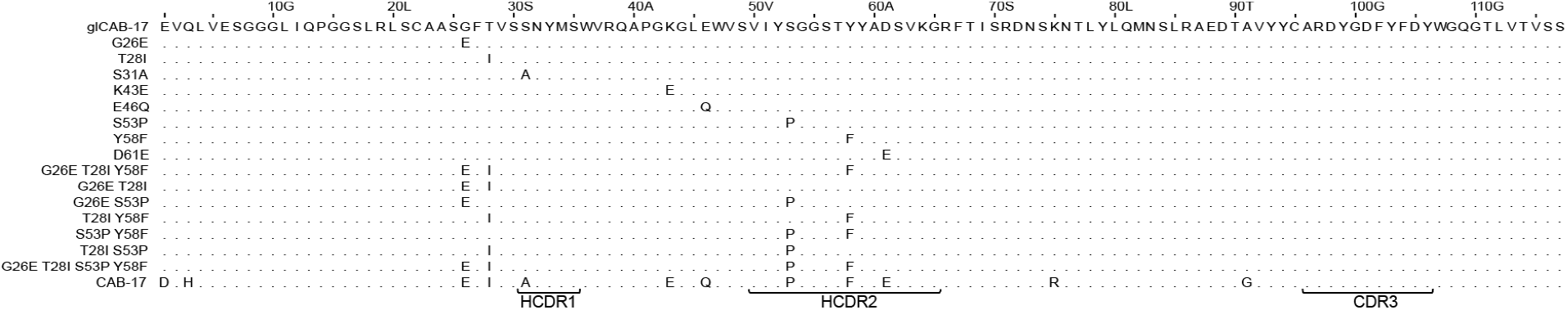
Alignment of the key HC SHMs introduced into glCAB-A17. Related to Fig. 2.

**Fig. S4.**
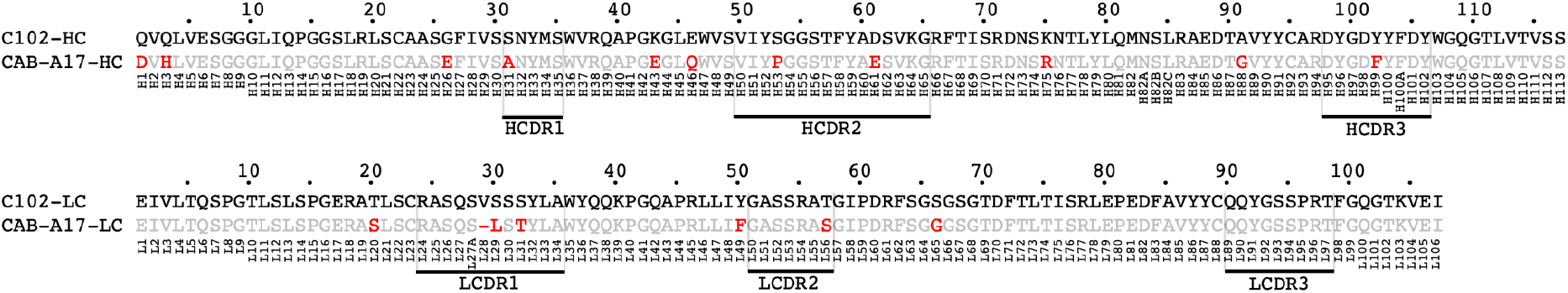
Alignment showing differences between CAB-A17 and C102.

**Fig. S5.**
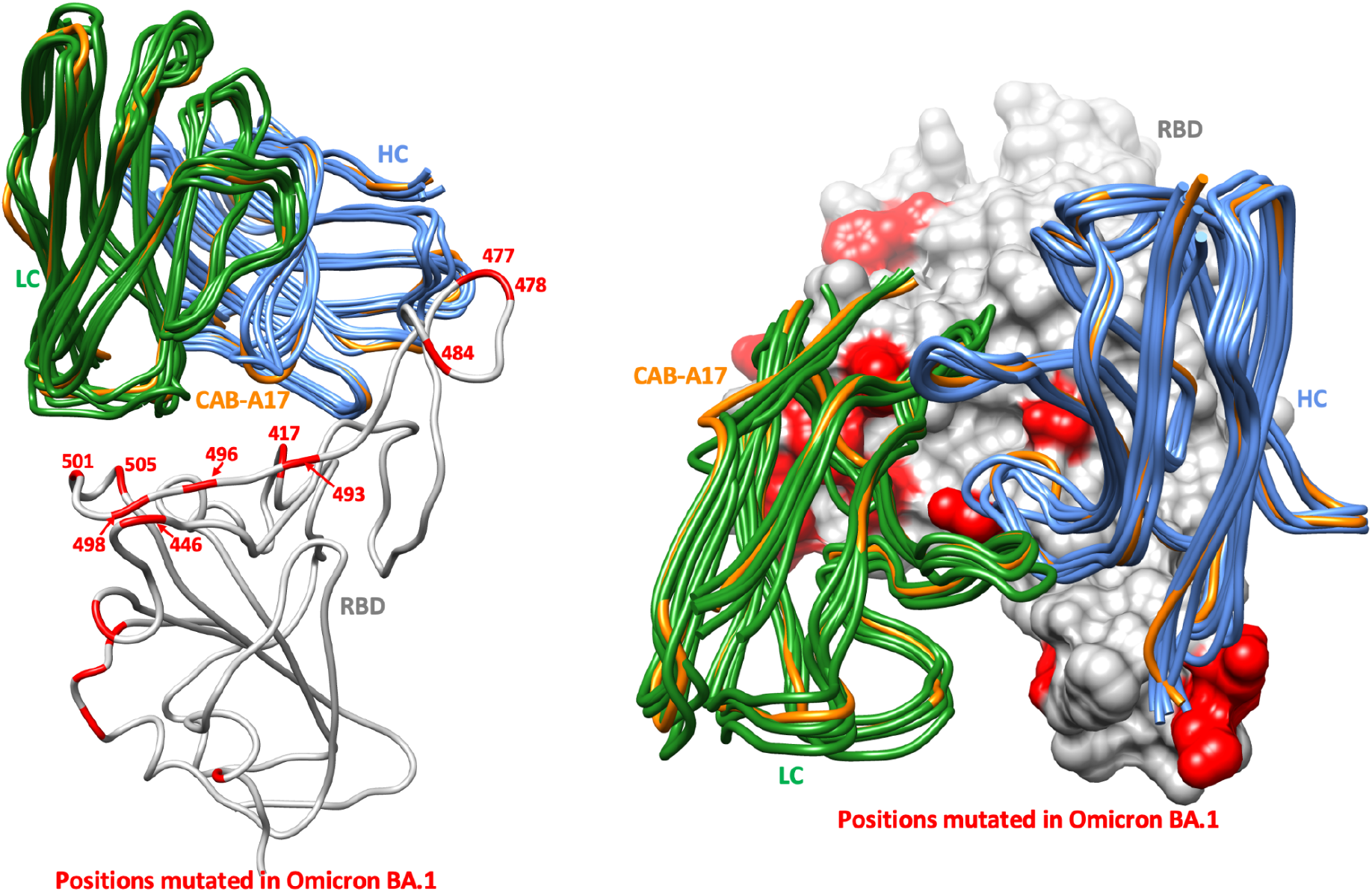
Superposition of CAB-A17 (orange) with the 5 mAbs in figure 1A.

**Table S1.** Hydrogen bond table CAB-A17 – Omicron.

**Heavy chain CAB-A17 hydrogen bonds – Omicron RBD**
| CAB-A17 HC residue, moiety | Distance (Å) | Omicron residue<br>(mutations in **red**),<br>moiety |
| --- | --- | --- |
| I28 N | 3.16 | A475 O |
| N32 ND2 | 2.74 | A475 O |
| Y33 OH | 3.68 | **N417** ND2 |
| Y33 OH | 2.28 | L455 O |
| G54 N | 2.96 | Y421 OH |
| S56 OG | 2.84 | T415 OG1 |
| S56 OG | 2.85 | D420 OD2 |
| R97 NH1 | 3.44 | N487 OD1 |
| R97 NH2 | 3.00 | Y489 OH |
| E26 O | 3.08 | N487 ND2 |
| A31 O | 2.14 | Y473 OH |
| G54 O | 2.51 | N460 ND2 |

**Light chain CAB-A17 hydrogen bonds – Omicron RBD**
| CAB-A17 LC residue, moiety | Distance (Å) | Omicron residue<br>(mutations in **red**),<br>moiety |
| --- | --- | --- |
| Y32 OH | 3.68 | **S496** OG |
| T31 OG1 | 3.74 | **Y501** OH |
| S30 OG | 2.95 | H505 ND1 |
| G92 O | 3.14 | R403 NH2 |

**Table S2.** Cryo-EM data collection, refinement and validation statistics.

|  | Omicron spike +<br>Fab CAB-A17<br>(EMDB-xxxx)<br>(PDB xxxx) | Omicron +<br>Fab CAB-A17<br>(EMDB-xxxx)<br>(PDB xxxx) |
| --- | --- | --- |
| **Data collection and processing** |  |  |
| Magnification (nominal) | 165,000 |  |
| Voltage (kV) | 300 |  |
| Electron exposure (e-/Å<sup>2</sup>) | 54.6 |  |
| Underfocus range (µm) | 0.3–2 |  |
| Pixel size (Å) | 0.505 |  |
| Symmetry imposed | C1 |  |
| Initial particle images (no.) | 1,826,695 |  |
| Final particle images (no.) | 398,386 | 57,905 |
| Map resolution (Å) | 2.56 | 3.62 |
| FSC threshold | 0.143 | 0.143 |
| **Refinement** |  |  |
| Initial model used (PDB code) | 7A29 | 7A29+7K8M |
| **Map sharpening *B* factor (Å<sup>2</sup>)** | **107.1** | **142.0** |
| **Model composition** |  |  |
| Non-hydrogen atoms | 30930 | 4797 |
| Protein residues | 3873 | 619 |
| Ligands | 52 NAG | - |
| ***B* factors (Å<sup>2</sup>)** |  |  |
| Protein | 82.4 | 84.9 |
| Ligand | 103.9 | - |
| **R.m.s. deviations** |  |  |
| Bond lengths (Å) | 0.003 | 0.002 |
| Bond angles (°) | 0.678 | 0.557 |
| **Validation** |  |  |
| MolProbity score | 1.62 | 1.29 |
| Clashscore | 3.29 | 4.99 |
| Poor rotamers (%) | 0.67 | 0.19 |
| **Ramachandran plot** |  |  |
| Favored (%) | 98.24 | 97.87 |
| Allowed (%) | 1.76 | 2.13 |
| Disallowed (%) | - | - |

